# Bacterial motility patterns vary smoothly with spatial confinement and disorder

**DOI:** 10.1101/2024.09.29.615714

**Authors:** Haibei Zhang, Miles T. Wetherington, Hungtang Ko, Cody E. FitzGerald, Leone V. Luzzatto, István A. Kovács, Edwin M. Munro, Jasmine A. Nirody

## Abstract

In unconfined environments, bacterial motility patterns are an explicit expression of the internal states of the cell. Bacteria operating a run-and-tumble behavioral program swim forward when in a ‘run’ state, and are stalled in place when in a reorienting ‘tumble’ state. However, in natural environments, motility dynamics often represent a convolution of bacterial behavior and environmental constraints. Recent investigations showed that *Escherichia coli* swimming through highly confined porous media exhibit extended periods of ‘trapping’ punctuated by forward ‘hops’, a seemingly drastic restructuring of run-and-tumble behavior. We introduce a microfluidic device to systematically explore bacterial movement in a range of spatially structured environments, bridging the extremes of unconfined and highly confined conditions. We observe that trajectories reflecting unconstrained expression of run-and-tumble behavior and those reflecting ‘hop-and-trap’ dynamics coexist in all structured environments considered, with ensemble dynamics transitioning smoothly between these two extremes. We present a unifying ‘swim-and-stall’ framework to characterize this continuum of observed motility patterns and demonstrate that bacteria employing a consistent set of behavioral rules can present motility patterns that smoothly transition between the two extremes. Our results indicate that the control program underlying run-and-tumble motility is robust to changes in the environment, allowing flagellated bacteria to navigate and adapt to a diverse range of complex, dynamic habitats using the same set of behavioral rules.

---

A diverse group of flagellated bacteria, including *E. coli* and *Salmonella*, use a motility strategy called ‘run-and-tumble’. Briefly, bacteria swim forward when their flagellar filaments come together to form a rotating bundle (‘runs’), and reorient themselves when this bundle comes apart (‘tumbles’). In free solution, bacteria can execute this behavioral program with no environmental constraints. This produces trajectories that resemble a biased random walk, with ballistic runs in a persistently directed path punctuated by short tumbles that randomly reset the direction of motion. [1, 2, 3, 4, 5, 6].

Recent work showed that *E. coli* in tightly packed hydrogels display motility patterns strikingly different from those seen in unconfined environments. In highly confined environments, bacteria remain ‘trapped’ for long periods until they are able to reorient themselves to ‘hop’ through an opening between pores [7, 8]. What environmental factors contribute to the differences in motility dynamics? How do motility patterns transition between those observed in these two extreme cases? Here, we introduce a microfluidic device to rigorously characterize the impact of environmental structure on *E. coli* motility patterns, by systematically varying two parameters: confinement, a proxy for obstacle density, and disorder, a proxy for obstacle randomness (Fig. 1).

**Figure 1:**
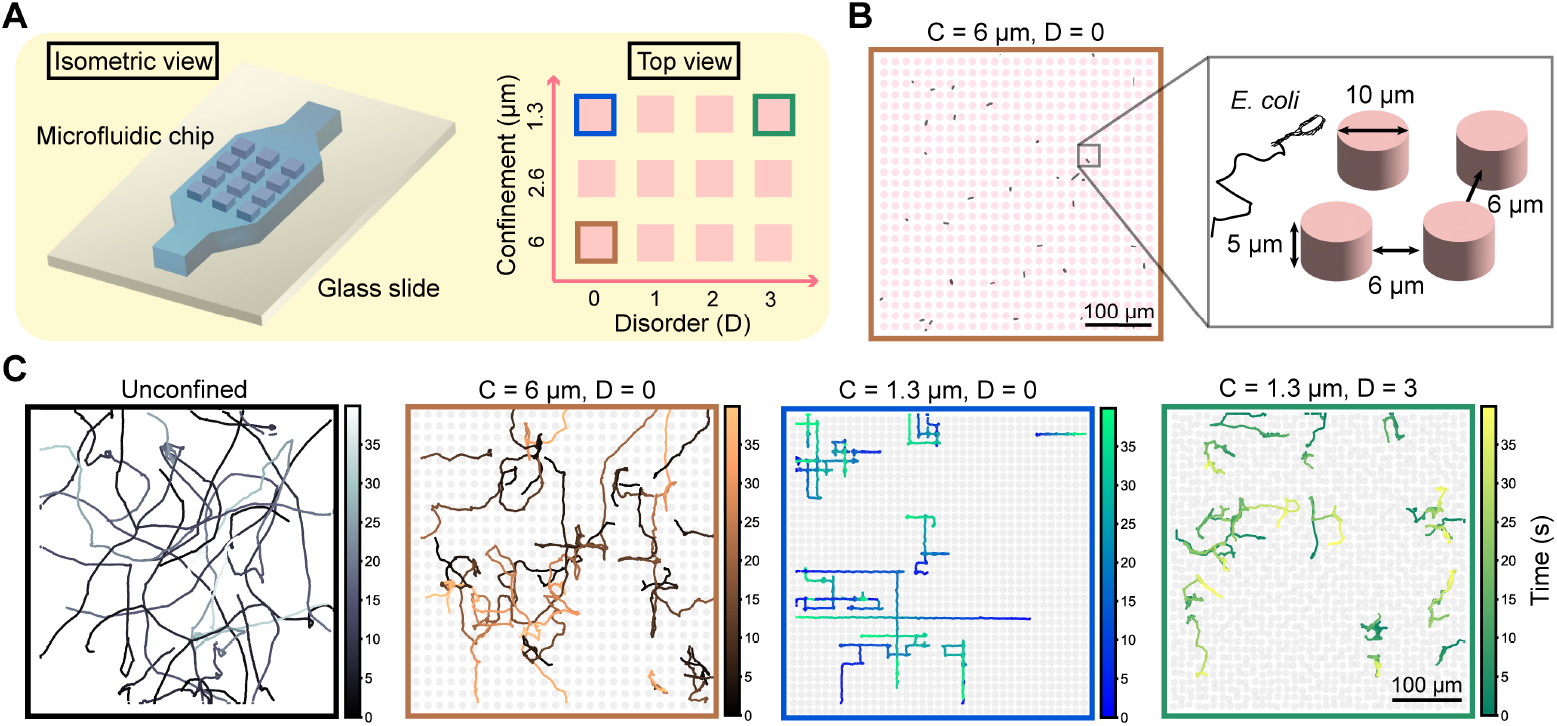
A microfluidic device exposing bacteria to environments of varying levels of confinement and disorder. (**A**) Schematic of our experimental device in isometric and top views. The PDMS microfluidic chip comprises twelve regions, represented as pink rectangles. Each region exposes bacteria to a different level of confinement *C* and disorder *D*. (**B**) An example environment with *C* = 6 *µ*m, *D* = 0. Inset shows a schematic of pillar spacing with respect to a swimming *E. coli* cell. (**C**) Representative tracked trajectories for unconfined and three structured regions (boxed in corresponding colors in **A**). Twenty randomly selected trajectories are shown for each region; color gradient along the track indicates time lapse (darker to lighter coloring denotes time from start to end of the trajectory). Cells are tracked using ImageJ plugin ‘TrackMate’. All regions are of size 400*µ*m x 400*µ*m; the unconfined region has no pillars.

We demonstrate that ensemble-level dynamics smoothly transition from those characteristic of unconstrained run-and-tumble to those characteristic of hop-and-trap as confinement and disorder increase. Trajectories consistent with both extremes coexist in nearly all structured regions considered. This suggests that bacteria display a continuum of motility patterns that cannot be characterized by a set of discrete locomotive modes. We present a unifying ‘swim-and-stall’ framework to characterize observed patterns across this continuum, where ‘swims’ are defined as periods of directed forward movement (e.g., ‘runs’ or ‘hops’) and ‘stalls’ are defined as the non-processive periods between adjacent swims (e.g., ‘tumbles’ or ‘traps’). Transitions along the motility continuum are driven by changes in both swim and stall duration. We quantify the contributions of these changes through the variation of two tuning parameters: *apparent tumble frequency*, the average rate at which swims terminate controls the swim duration; and *tumble failure probability*, the likelihood that a bacterium fails to initiate a period of straight swimmming after a reorientation event, controls the stall duration.

We further explore whether transitions in motility dynamics observed across conditions are driven by environmental constraints, changes in the internal states of individual bacteria, or some combination of the two. A swim period can be terminated through a collision with an obstacle (environmental constraint) or through the initiation of a tumble by a bacterium (change in internal state). Similarly, a stall period can be extended by a bacterium keeping its flagellar bundle splayed apart (internal state) or through a tight spot preventing it from swimming forward (environmental constraint). Migrating populations of bacteria have been shown to tune their swimming behavior in response to physical and chemical stimuli from their environment [2, 9, 10]. Do bacteria make similar adjustments to their underlying behavioral program in response to mechanical stimuli from their environment? We show evidence that bacteria in unconfined and highly confined environments are operating the same run-and-tumble motility program, just in different settings – that is, no change in bacterial behavior is needed to transition between motility ‘modes’ at opposing ends of the continuum. We provide context for how quantifiable environmental constraints – confinement and disorder – impact our proposed tuning parameters and show that these are sufficient to explain the changes in motility patterns we observe across environments. Our results indicate that flagellated bacterial species executing a run-and-tumble motility program can efficiently navigate a range of constrained and heterogeneous environments using a universal set of behavioral rules.

## Results

### A microfluidic system to investigate bacterial motility in spatially structured environments

We construct a microfluidic device to quantify how mechanical challenges posed by natural environments [11, 12, 13, 14] affect *E. coli* motility patterns using two parameters: (1) confinement, a measure of the density of obstacles in a region; and (2) disorder, the level of deviation from a perfect grid. Our device comprises twelve confined regions and one unconfined region, each of size 400 *µ*m x 400 *µ*m. Confined regions contain pillars of 5 *µ*m in height and 10 *µ*m in diameter in varying arrangements (Fig. 1A, 1B); the unconfined region contains no pillars.

Three levels of confinement *C* are considered: *C* = 6 *µ*m, 2.6 *µ*m, and 1.3 *µ*m, where *C* is the mean spacing between pillars. The chosen values range from the length of a flagellar filament (∼ 7µm) at the lowest level of confinement to the length of the cell body (∼ 2µm) at the highest [15, 16]. For levels of disorder D, the center of each pillar is shifted away from the grid position by a value sampled from a uniform distribution with range 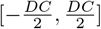; *D* = 0 corresponds to a perfect grid arrangement (Fig. 1A). For example, for *C* = 6 *µ*m, *D* = 0 region, pillars are horizontally and vertically spaced 6 *µ*m apart (Fig. 1B); whereas for *C* = 1.3 *µ*m, *D* = 3, pillars are initially spaced 1.3 *µ*m apart and their positions are displaced by 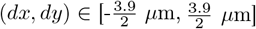. Fluorescently labeled *E. coli* in a nutrient-rich medium are flowed into the device for observation (Supplementary Movies 1-3).

Each trial records bacterial movement for forty seconds at a time resolution of 0.05 s (20 frames per second). Randomly sampled trajectories from three structured conditions and in unconfined environments are shown in Fig. 1C (n = 20 trajectories from each region). We observe qualitative differences in bacterial trajectories in structured environments and unconfined media (Fig. 1C). For instance, trajectories tend to be directed vertically or horizontally in confined, ordered regions, indicating that bacteria move along rows and columns of pillars rather than performing a uniform exploration of the space as in unconfined environments. Accordingly, we find that trajectories become more tortuous as disorder increases (Fig. 1C and Supplementary Fig. 1). In the following sections, we quantitatively analyze these observations.

### Swim-and-stall: a unifying framework across environments

Historically, bacterial motility states (run vs. tumble) correspond to flagellar orientations (bundled vs. unbundled) [2]. In free solution, there is no constraint to a bacterium expressing its internal state. However, the one-to-one mapping between internal and observed states can break down in complex environments [7, 8]. Accordingly, a consistent framework for distinguishing between a bacterium’s observed behavior and its internal state is crucial for making meaningful comparisons across conditions.

We introduce a unifying framework that classifies observed bacterial motion into two states: ‘swims’, periods of directed movement (corresponding to ‘runs’ or ‘hops’), and ‘stalls’, non-processive periods (corresponding to ‘tumbles’ or ‘traps’). These designations describe the observed dynamics of bacteria based on their center-of-mass (COM) trajectories, without direct knowledge of flagellar motor dynamics. In unconfined environments, observed and internal states align — all swims are runs, and all stalls are tumbles. However, in confined environments, obstacles can prematurely terminate a swim due to bacterial collisions with obstacles or extend a stall by physically trapping a cell (Fig. 2A).

**Figure 2:**
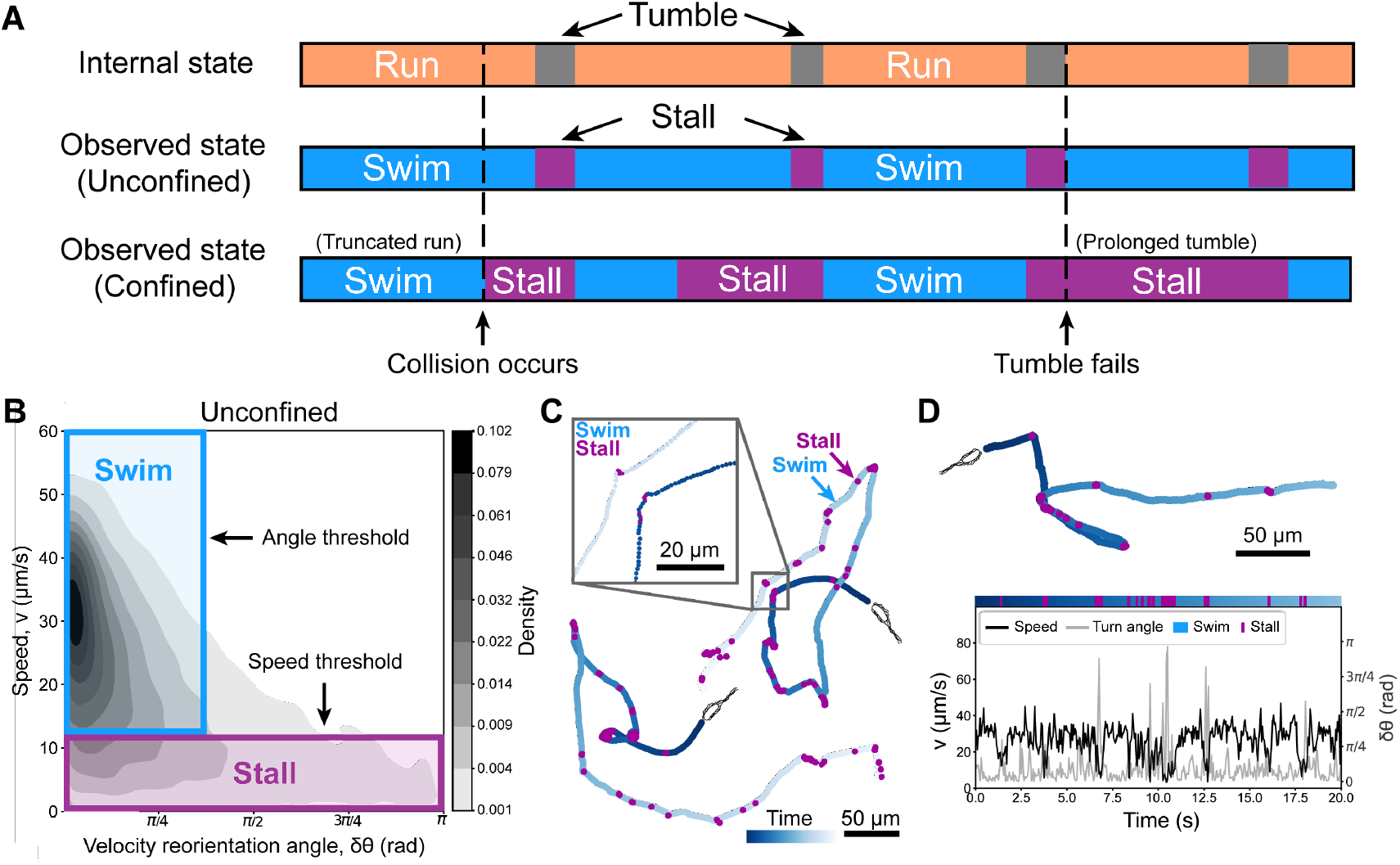
Classification of bacterial trajectories. (**A**) Schematic demonstrating how bacteria employing the same run-and-tumble behavioral program (internal state, top) can display differences in observed state in unconfined regions (middle) and confined regions (bottom). First, collisions with obstacles in confined regions can result in swim periods that conclude before a full run is executed (see the first arrow at the bottom bar). Second, confined environments can trap a bacteria in place at the conclusion of a tumble, resulting in a prolonged stall period that comprises multiple internal state transitions (see the second arrow at the bottom bar). (**B**) The bivariate kernel density estimate shows the distributions of bacterial speeds *v* and turn angles *δθ* in the unconfined region. Swims (boxed in blue) occur when *v* > 12.4 *µ*m/s (half the mean speed) and *δθ* < 3*π* rad. Stalls (boxed in purple) occur when the *v* < 12.4 *µ*m/s. (**C**) Two sample trajectories from bacteria in the unconfined region, with swims and stalls colored in blue and purple, respectively. (**D**) Twenty seconds of a sample trajectory in the unconfined region. Speed (black) and turn angle (grey) values for each frame determine the behavioral state (‘swim’ or ‘stall’) assigned to each frame. Color bar above shows discrete sequence of states, with frames assigned as swims shown in blue and stalls in purple.

Implementing a consistent discrete state mapping across conditions under this new framework allows us to quantify if and how motility patterns change in response to environmental structure. Swimming speeds in the unconfined region, 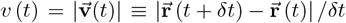, are narrowly distributed with mean speed ⟨*v*⟩ = 24.8 *µ*m/s, consistent with previously reported *E. coli* swimming speeds of 20-30 *µ*m/s [2, 17]. Velocity reorientation angles (turn angles), 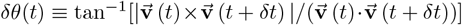, are broadly distributed from 0 to π (Fig. 2B). We denote a ‘swim’ period when a cell is moving at least half the mean unconfined speed, 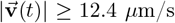, and its turn angle is less than or equal to π/3 radians (60*°*). We define ‘a stall’ period when a cell moves slower than half of the mean unconfined speed; turn angles for stall periods are widely distributed [7]. Speed and angle thresholds are shown in Fig. 2B; these criteria ensure that fewer than 5% of frames remain unclassified 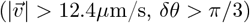, and results are robust to small threshold variations (Supplementary Fig. 2).

Representative trajectories in the unconfined region labeled according to these two states (swim and stall) are shown in Fig. 2C. Cells alternate between periods of directed motion and in-place reorientation. A sample timecourse of swimming speed and turn angle further illustrates this pattern: swims are characterized by high swimming speeds and low turn angles, and stalls with precipitous drops in speed and spikes in turn angle (Fig. 2D).

### Swim lengths decrease and stall bias increases with confinement

Using this swim-and-stall framework, we compare observed motility dynamics across different levels of confinement and disorder. For each condition, we compute the swim time ⟨*T*_swim_⟩ and the mean stall time ⟨*T*_stall_⟩ of all tracked cells (Fig. 3A, 3B). The mean swim time is highest in the unconfined region (⟨*T*_swim_⟩ = 0.77 s) and lowest in the most confined region (⟨*T*_*swim*_⟩ = 0.25 s for *C* = 1.3 *µ*m, *D* = 0). The swim time distribution for the unconfined is significantly different from all other conditions (p < 0.0001, Supplementary Fig. 6). The level of disorder does not significantly affect ⟨*T*_swim_⟩.

**Figure 3:**
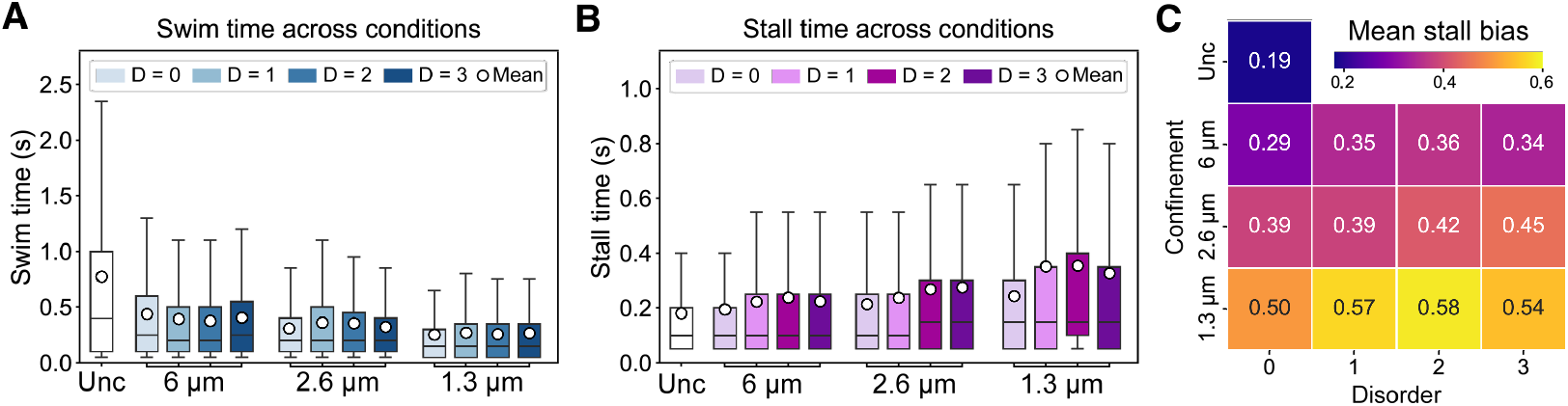
Dynamics of swims and stalls. (**A**) Swim time and (**B**) stall time distributions across experimental conditions shown in boxplots. All distributions are log normal with positive skew. The lines inside the boxes indicate the median values and white circles mark the mean values. (**C**) The heatmap of mean stall bias across experimental conditions of confinement *C* and disorder D. Mean stall bias is defined as the average stall time proportion over all tracked bacteria in a given region. The label ‘unc’ represents the unconfined region.

Stall times increase with both confinement and disorder; additionally, the variance in stall times increases in disordered regions (white circles in Fig. 3B). Stall times in all structured environments are significantly different from those in unconfined environments (p < 0.05, Supplementary Fig 7). We hypothesize that prolonged stalls comprise multiple state transitions that cannot be resolved experimentally (Fig. 2A, 3B). For instance, several reorientations may be required for bacteria to move past dead ends created by disordered pillar array geometries. We also see a wider distribution of the turning angle in these regions (Supplementary Fig. 3).

The mean stall (observed tumble) bias, ⟨*P*_stall_⟩, is the fraction of time at which cells stall on average, and n and is defined as 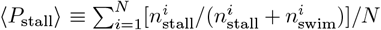 where 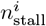 and 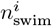 are the number of frames assigned to stall or swim states of the ith cell, respectively, and N is the total number of tracked bacteria. ⟨*P*_stall_⟩ is much higher in structured regions as compared to unconfined media (Fig. 3C), supporting the hypothesis that cells show shorter swim durations when they encounter physical obstacles (i.e., have a higher apparent tumble frequency). The increase in stall bias observed in structured environments likely arises from changes in both swim and stall duration.

### Characteristic patterns coexist in complex structured environments

In free solution, bacteria express their internal states with no constraints: bacteria operating a run-and-tumble behavior show ballistic swims punctuated by short reorienting stall periods [2]. These dynamics are captured clearly by examining the mean squared displacements, *MSD*(*t*) = ⟨(*r* (*t* + *τ*) − *r* (*t*))^2^⟩. The ensemble-averaged MSD for run-and-tumble behavior in unconfined environments varies quadratically (*ν* = 2) for short lag times *τ*, reflecting ballistic dynamics during runs (swims), and linearly (*ν* = 1) at long lag times, reflecting long-term diffusive dynamics. The crossover time between these two regimes *τ*_*c*_ corresponds to the mean duration of a swim period (Fig. 4A). Hop-and-trap behavior deviates from that of unconstrained run-and-tumble both at short and intermediate timescales. First, at short time lags, bacteria in highly confined media display superdiffusive (1.5 < *ν* < 2), rather than ballistic, dynamics. Second, at intermediate time lags, bacteria display transient subdiffusive (*ν* < 1) dynamics; these give way to diffusive behavior at longer timescales as in unconstrained run-and-tumble.

**Figure 4:**
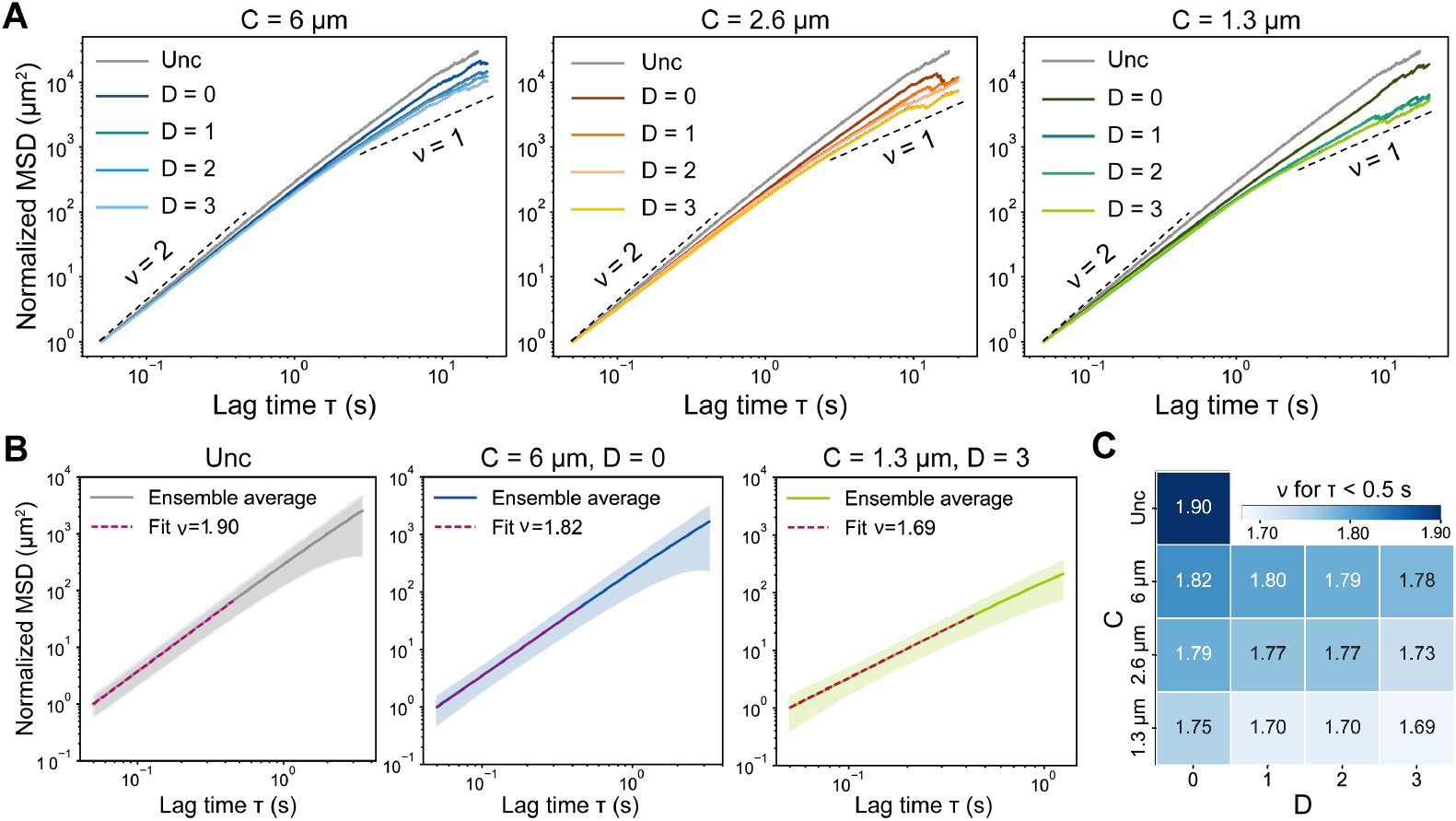
Long-range dynamics across varying levels of confinement and disorder. (**A**) Normalized ensemble-averaged mean square displacements (eMSDs) for *C* = 6 *µ*m (left), *C* = 2.6 *µ*m (center), and *C* = 1.3 *µ*m (right). Each plot depicts eMSD curves for varying levels of disorder D. Dashed lines show ∼ *t*^*ν*^ for ballistic (*ν* = 2) and diffusive motion (*ν* = 1). (**B**) Curves for eMSDs at early lag times for unconfined (left), *C* = 6 *µ*m; *D* = 0 (center), and *C* = 1.3 *µ*m, *D* = 3 (right). Dashed pink lines show fits estimating *ν* for *τ* < 0.5 s. The error bands indicate ± one standard deviation. (**C**) Fitted values of *ν* for *τ* < 0.5 s across confinement *C* and disorder D.

We observe bacterial trajectories consistent with both extremes in all structured environments considered. Close examination of individual MSDs reveals a high level of variance in structured regions, with the proportion of trajectories displaying ballistic dynamics decreasing as confinement and disorder increase (Supplementary Fig. 3). We find superdiffusive behavior 1.5 < *ν* < 2 at short time lags 0 < τ < 0.5 for all structured regions (Fig. 4B). The value of *ν* for short time lags decreases as confinement and disorder increase, and ensemble dynamics approach those consistent with the hop-and-trap model (*ν* ≈ 1.5, Fig. 4C) [7]. Regions that are both highly confined and highly disordered show ensemble dynamics closest to those expected from hop-and-trap (*ν* = 1.69 for *C* = 1.3 *µ*m, *D* = 3).

We similarly observe trajectories with transient subdiffusive behavior at all intermediate time lags, with the proportion of such trajectories increasing with both confinement and disorder (Supplementary Figs. 4 and 5). However, this trend can easily be masked when the variance in dynamics is large, as is the case at intermediate levels of confinement. This is consistent with previous observations that ensemble-averaged values of *ν* remain close to 1 until packing is very dense in porous media [7].

**Figure 5:**
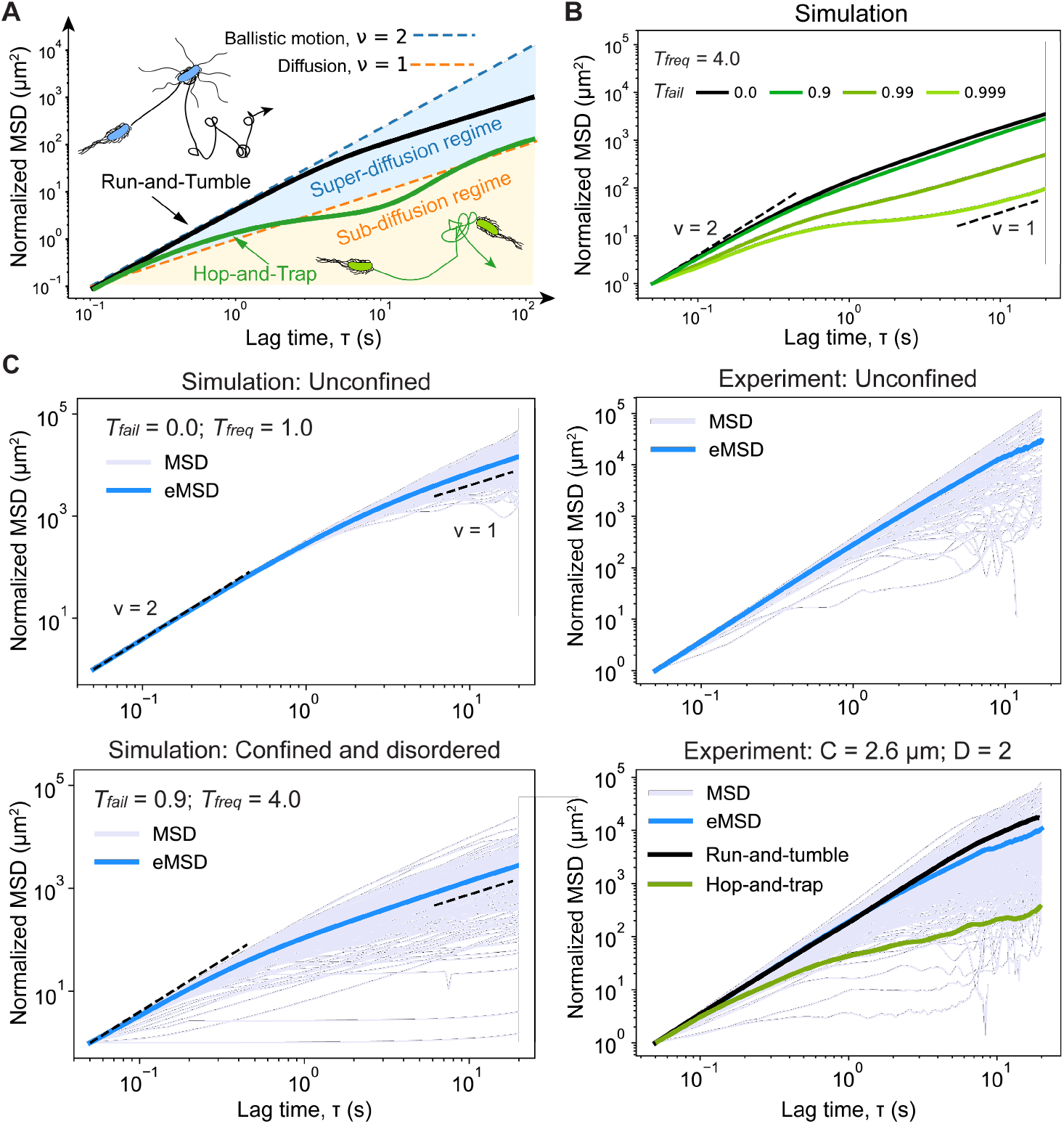
Characteristic motility patterns coexist in structured environments. (**A**) Schematic of characteristic MSD curves for observed dynamics for bacteria employing unconstrained run-and-tumble behavior and observed dynamics under the hop-and-trap model. (**B**) Normalized ensemble-averaged MSD plot of ensemble dynamics with *T*_freq_ = 0 and varying tumble failure, *T*_fail_. (**C**) Time-averaged MSDs (in lavender) and ensemble-averaged MSDs (in blue) from simulations and experiments. Simulation parameters are chosen according to those observed in the unconfined region, *T*_fail_ = 0.0 and *T*_freq_ = 1.0 (top left), and in a confined and disordered region, *T*_fail_ = 0.99 and *T*_freq_ = 4.0 (bottom left). Results are shown from the corresponding experiments in the unconfined region (top right) and *C* = 2.6 *µ*m, *D* = 2 (bottom right), respectively.

### Both swim and stall duration tune transitions between motility patterns

Our experiments show that environmental structure has two major impacts on bacterial motility patterns. First, swim periods are shortened as cells collide with obstacles. Second, stall periods are lengthened due to cells becoming ‘trapped’ by tight, tortuous pore spaces (Fig. 5). To untangle the effects of these changes on observed ensemble dynamics, we define two parameters: (1) apparent tumble frequency *T*_freq_ controls the duration of swim periods; (2) tumble failure probability *T*_fail_ controls the duration of stall periods.

Swim duration tunes the length of time that ballistic dynamics are observed; shorter swims (e.g., due to collisions with obstacles) result in a lower crossover time between ballistic and diffusive dynamics. Bacteria implementing run-and-tumble behavior in unconfined media show low apparent tumble frequencies, with swim duration far exceeding stall duration (Fig. 3B, Supplementary Fig. 10). In highly confined environments, bacteria show far shorter swim (‘hop’) durations [7, 8]. At such high apparent tumble frequencies (⟨*T*_swim_⟩ = 0.12 s when *T*_freq_ = 8.0), swim periods may be so short that the ballistic region is difficult to resolve at experimental resolution (dt = 0.05 s). This results in the appearance of superdiffusive behavior at small time lags, as is noted in hop-and-trap dynamics (Supplementary Fig. 11, top left panel).

On the other hand, the appearance of subdiffusive dynamics at intermediate time lags requires lengthening stall duration [7]; subdiffusive dynamics are not observed even at the highest apparent tumble frequencies when *T*_fail_ = 0 (Supplementary Fig. 11). Transient subdiffusion has been observed in systems with long trapping periods within heterogeneous environments [18, 19, 20, 21]; in bacterial systems, we might expect such subdiffusive behavior because cells are ‘trapped’ in place during prolonged stall periods comprising multiple transitions when tumble failure probability is high (Fig. 2A). In our simulations, we observe an increase in the proportion of trajectories displaying transient subdiffusive behavior as *T*_fail_ increases and cells are more likely to show longer stall (prolonged tumble) periods (Fig. 5C, left column; Supplementary Fig. 12). This is in agreement with our experimental results, which show high variance in the individual MSDs as environmental confinement and disorder are introduced (Fig. 5C, right column; Supplementary Fig. 4). Additionally, transient subdiffusion at the ensemble level appears only at high values of *T*_fail_ (Fig. 5B; Supplementary Fig. 11). This supports the absence of ensemble-level subdiffusive behavior in our experiments (Fig. 4), and aligns with findings that transient subdiffusion emerges only when stall durations significantly exceed swim durations [7].

### Environmental structure affects observed, but not internal, states

While environmental structure clearly affects observed motility patterns, it remains unclear whether it also affects transitions between *internal states*. That is, do individual bacteria adapt their motility strategy (e.g., vary the rate at which frequency at which tumbles are attempted or concluded) to improve performance as they receive mechanical feedback from their environment? Or are the changes in apparent tumble frequency and tumble failure probability we characterize above mainly driven by environmental constraints (e.g., collisions and trapping) with the underlying run-and-tumble behavioral program remaining consistent across conditions (Fig. 2A)?

To address these questions, we calculate the distribution of conditioned swim lengths that approximates the swim lengths expected if bacteria do not change their underlying behavioral program – that is, if apparent tumble frequency only changes due to collisions with an obstacle. This distribution is computed by considering the distribution of unconstrained swim lengths and the distribution of chord lengths – the length of free straight paths starting from randomly chosen points within a given pillar geometry – and the minimum of the two random lengths is taken [22, 23] (Fig. 6). The chord length distribution describes the straight paths available to bacteria given a randomly chosen initial position and a random direction (Supplementary Fig. 8).

**Figure 6:**
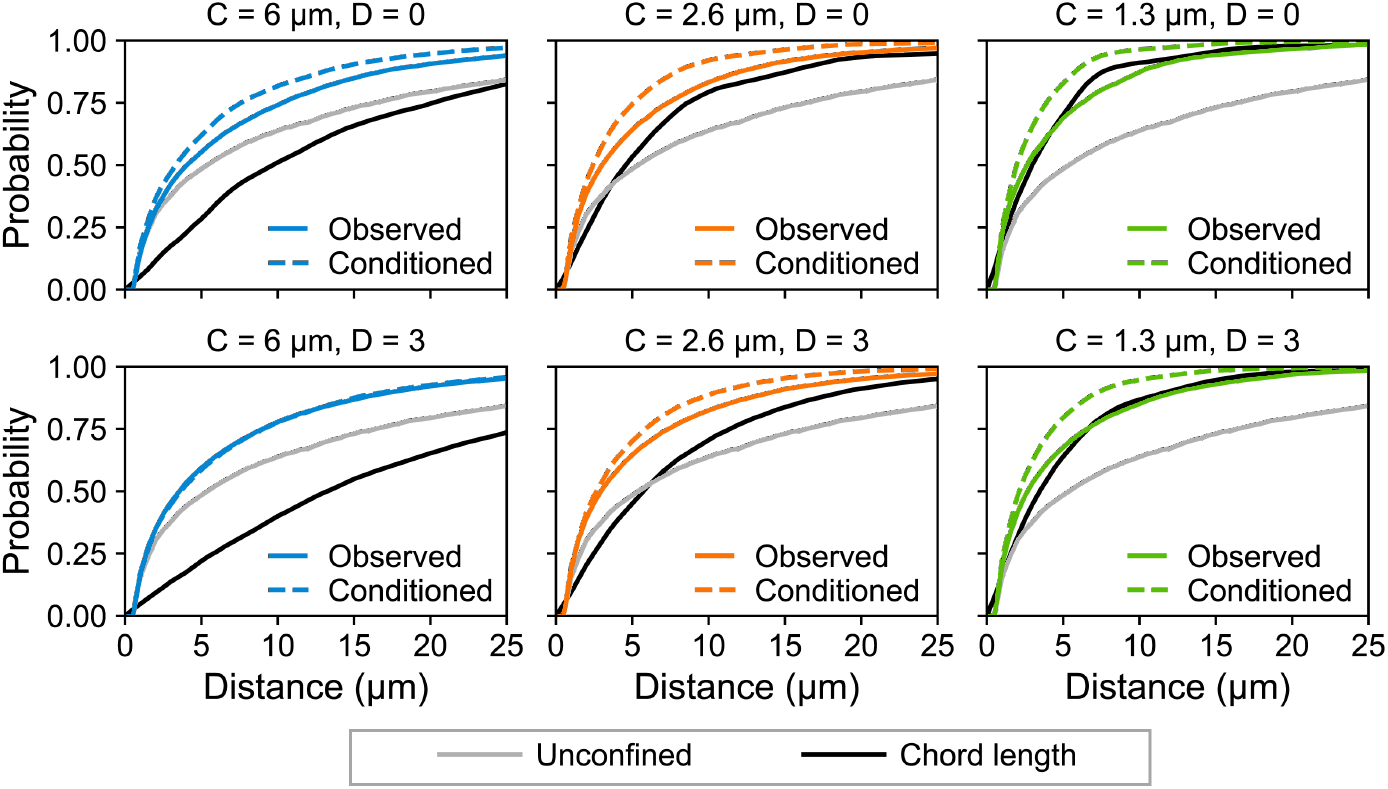
Observed and conditioned swim lengths for bacteria employing swim-and-stall behavior in confined and disordered environments. Cumulative distribution function (CDF) plots comparing observed (labeled in blue, orange, or green) and conditioned swim lengths (corresponding dashed lines) for *E. coli* across levels of confinement (increasing from left to right) and disorder (increasing from top to bottom). The CDFs for unconfined swim lengths (gray) and chord lengths (black) are also shown in each panel. The conditioned distribution corresponds to the minimum of the unconfined swim length and the chord lengths; this assumes that changes in swim lengths arise solely from environmental constraints, with no change in internal state transitions across conditions.

We find that observed swim lengths match the conditioned swim lengths well in disordered environments with low to intermediate confinement (e.g., for *C* = 6 *µ*m, *D* = 3 and *C* = 2.6 *µ*m, *D* = 3, *R*^2^ = 0.998 and 0.983, respectively). This strongly indicates that there is no change to the underlying behavioral program of bacteria as they are navigating these environments. In highly confined environments, the observed distribution of swim lengths aligns most closely with the chord length distribution (e.g., for *C* = 1.3 *µ*m, *D* = 0 and *C* = 1.3 *µ*m, *D* = 3, *R*^2^ = 0.978 and 0.963, respectively), suggesting that the availability of free straight paths is mainly responsible for constraining swim lengths in this regime. This is again a strong indicator that the geometry of the environment, rather than bacterial behavior, is the primary feature driving changes in observed motility patterns. Similarly, in environments with intermediate confinement and high order (C = 2.6 *µ*m, *D* = 0), we find that the observed distribution matches our prediction closely for short swim lengths but matches the chord length distribution for longer swim lengths (Fig. 6).

## Discussion

Environmental structure can interfere with the unconstrained execution of a bacterium’s behavioral program. We present a unifying ‘swim-and-stall’ framework for characterizing *E. coli* motility patterns across a diversity of environments, from free solution to highly confined and disordered media (Fig. 7A). Within this framework, ‘swims’ are periods of directed forward movement (whether they are concluded through an external collision or internal bacterial biochemistry) and ‘stalls’ are non-processive periods between adjacent swims (whether they correspond to ‘trapped’ or ‘tumbling’ cells). We show that observed motility patterns vary smoothly with both confinement and disorder and that chanages in swim and stall duration impact ensemble-level dynamics at short and intermediate timescales, respectively.

**Figure 7:**
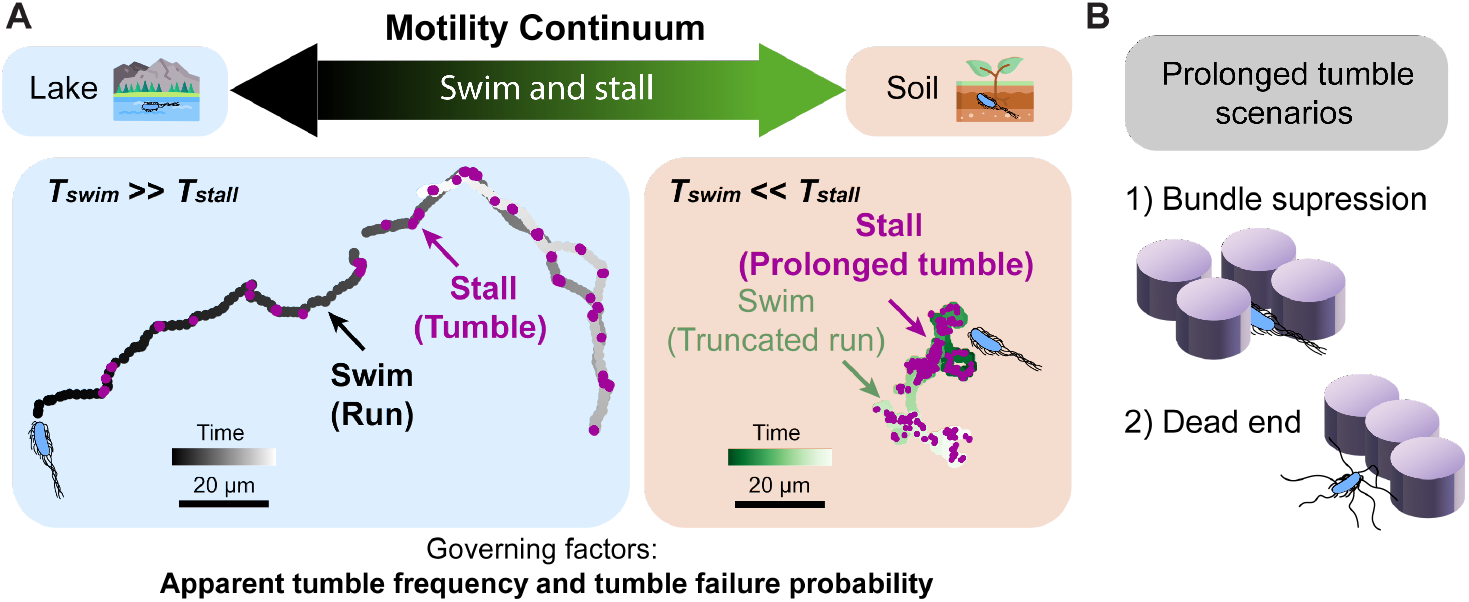
Bacteria executing run-and-tumble behavior display a continuum of motility patterns tuned by duration of swim and stall periods. (**A**) Two highlighted trajectories are typical of run-and-tumble and hop-and-trap models corresponding to MSD curves in Fig. 5C with *C* = 2.6 *µ*m; *D* = 2. (**B**) Two scenarios where stall periods might be prolonged due to environmental factors: 1) the flagellar bundle is suppressed in confined environments; 2) dead end blocks forward movement after reorientation in disordered environments.

As observed motility dynamics often represent an entanglement of bacterial behavior and environmental constraint, we investigate what drives changes in these motility parameters across conditions. We compute the distribution of swim lengths observed in the unconfined region conditioned on the geometry of each structured environment considered to investigate whether *E. coli* modify their behavior in response to changes in environmental structure (Fig. 6). At low and intermediate levels of confinement, we find that our predicted distribution matches well with observed swim lengths. At the highest level of confinement and disorder, we find that trajectories tend to mirror environmental geometry, with the distribution of observed swim lengths closely matching the distribution of possible straight paths. This suggests that, in extreme environments, bacterial ‘decisions’ are largely overruled by environmental constraints, and that observed motility patterns in such environments may be independent of changes in the underlying run-and-tumble behavioral program. For instance, a ‘stalled’ bacterium may not able to display forward movement (i.e., enter a swim period) after concluding a tumble (*T*_fail_ > 0). In this case, the internal state and the observed state of a cell during a stall period deviate – that is, a single resolvable stall period may actually comprise multiple internal state transitions due to ‘failed tumbles’ (Fig. 2A). In the case of a single failed reorientation, a stall would be composed of a tumble (resulting in a failed reorientation), a non-progressive run, and a second tumble (resulting in a successful reorientation). Prolonged stall periods may include multiple failed transitions between actual tumbles and non-progressive runs. Tumble failures occur in two environmentally-driven situations (Fig. 7B): (1) tight spacing forces the flagellar bundle to stay together, suppressing reorientation (high confinement) or (2) a reorientation does not allow the subsequent run to be progressive because of an immediate collision with another obstacle within a ‘dead end’ cluster (high disorder) [7].

Deviations between the conditioned and observed distributions in highly confined and highly ordered conditions likely arise from our assumptions that (1) all positions are equally likely to be the starting point of a swim and (2) all directions are equally likely to be chosen at the start of a swim. In highly confined environments, the presence of dead ends might lead to trapping, making the density of trajectories highly non-uniform. In highly ordered environments, shallow collision angles can promote alignment with the surface and narrow spacing likely restricts cell reorientation during a tumble.

Our results indicate that *E. coli* do not modify swimming behavior in response to mechanical stimuli on short to intermediate timescales (seconds to minutes). Therefore, observed changes in motility patterns arise from bacterial ‘decisions’ being thwarted by environmental constraints. An interesting direction for future work would be to explore if adaptation of run-and-tumble behavior to optimize motility is observed on longer timescales in given environments, particularly in the context of chemical gradients.

*E. coli*, and other peritrichously flagellated bacterial species, are found in nearly every environment imaginable, and must contend with a wide range of heterogeneity and uncertainty in their surroundings [24, 14, 5] (Fig. 7). Other bacterial motility strategies such as ‘run-and-reverse’ – used primarily in marine bacteria like *Vibrio alginolyticus* and *Shewanella putrefaciens* – outshine run-and-tumble in specific contexts, such as adeptly responding to nutrient patches in the ocean by rapidly climbing chemical gradients [25, 26]. This behavior (and its variants, including run-reverse-flick and run-reverse-wrap) can also be adapted to avoid getting trapped in tight spaces [27, 28]. This is especially crucial for symbiotic species like *Vibrio fischeri*, as they must navigate to and through specific host tissues, often through highly confined and highly viscous regions [29, 30]. While run-reverse strategies seem to prioritize success in structurally constant environments, the run-and-tumble motility program must be more broadly successful [22, 28, 31, 32]. Indeed, when faced with complex or heterogeneous environments, *V. alginolyticus* grow lateral flagella and adopt a ‘generalist’ strategy more akin to run-and-tumble [33]. Our work underscores both how environment can shape our observation of a bacterium’s behavior, as well as how the range of environments a bacterium must successfully navigate can shape the evolution of its underlying behavioral program.

## Acknowledgments

We thank Tom Pennell for his guidance and advice throughout the fabrication process at the Cornell NanoScale Facility. We also thank Benjamin R. Epley, Erin Brandt, Peter J. Yunker and Juan E. Keymer for their insights and many thoughtful discussions.

H.Z. and J.A.N. acknowledge funding support by NSF-Simons National Institute for Theory and Mathematics in Biology, which is jointly supported by the U.S. National Science Foundation (Award 2235451) and the Simons Foundation (Award MP-TMPS-00005320). H.Z. is also grateful for her graduate program in Biophysical Sciences at the University of Chicago. M.T.W., H.K., C.E.F., and J.A.N. acknowledge support from The Santa Fe Institute and The James S. McDonnell Foundation Postdoctoral Fellowship Award in Complex Systems (M.T.W: https://doi.org/10.37717/2020-1543; H.K.: https://doi.org/10.37717/2021-3524; C.E.F.: https://doi.org/10.37717/2020-1591; J.A.N.: https://doi.org/10.37717/220020527). C.E.F. is supported by the NSF-Simons Center for Quantitative Biology at Northwestern University (NSF: 1764421 and Simons Foundation/SFARI 597491-RWC). J.A.N. and E.M.M. acknowledge support from the National Science Foundation through the Center for Living Systems (Grant # 2317138). This work was performed in part at the Cornell NanoScale Facility, a member of the National Nanotechnology Coordinated Infrastructure (NNCI), which is supported by the National Science Foundation (Grant NNCI-2025233).

## Author contributions

M.T.W., H.K., C.E.F., H.Z., and J.A.N. designed the research; M.T.W. performed experiments; H.Z. analyzed the data; J.A.N. performed simulations; L.V.L. performed chord length distribution calculations; H.Z. and J.A.N. wrote the manuscript; H.Z., M.T.W., H.K, C.E.F., L.V.L., I.A.K., E.M.M, and J.A.N. reviewed and edited the manuscript.

## Data and materials availability

Tracking data, analysis and simulation outputs have been deposited in Github [34].

