## Supplementary Information for "Bacterial motility patterns vary smoothly with spatial confinement and disorder"

### Supplementary Materials for
### Bacterial motility patterns vary smoothly with spatial confinement and disorder

Haibei Zhang[†], Miles T. Wetherington[†], Hungtang Ko, Cody E. FitzGerald,
Leone V. Luzzatto, István A. Kovács, Edwin M. Munro, Jasmine A. Nirody[*]

**This PDF file includes:**

Materials and Methods

Figures S1 to S11

Captions for Movies S1 to S3

**Other Supplementary Materials for this manuscript:**

Movies S1 to S3

### Materials and Methods

#### Strains and growth conditions

We grow a culture of wild-type *E. coli* (HCB33) with plasmids expressing red (mScarlet-I) fluorescent proteins on solid LB agar plates (LB Broth EZMix, Sigma-Aldrich + 1.5% Bacto Agar, MOLAR Chemicals). A single colony is subsequently inoculated in 3 mL LB medium (LB Broth EZMix, Sigma-Aldrich) with 50 $\mu$g/mL Ampicillin and Streptomycin for 16 h ± 30 min at 30 C° and 250 rpm. Overnight cultures are back-diluted 1:10,000 in the same medium; this culture has an OD600 of 0.5. 10 $\mu$L of this culture is gently mixed with 990 $\mu$L of fresh LB medium and antibiotics.

#### Microfluidic device fabrication and preparation

We design the pillar device using Klayout. We perform a direct write of the pillar device design on a 12.7-cm Cr mask plate (Heidelberg Mask Writer, DWL2000). We subsequently develop the patterns (Hamatech Mask Chrome Etch 1) and then place the mask plate for 2x10 minutes in a hot strip bath.

We prime a new Si wafer by spin coating (3000 rpm/1000 rps/30 s) a thin layer of MicroPrime MP-P20 (ShinEtsu MicroSi, Inc.) to promote adhesion for photoresist. Next, we spin coat Shipley 1813 photoresist on the wafer (using the same protocol) and immediately place it onto a 115 C° hotplate for 60 s. We perform surface analysis (Filmetrics) to check for surface irregularities. The outer (~1 cm) of photoresist is removed from the edge of the wafer with an Edge Bead Removal System. Using ABM's High Resolution Mask Aligner, we expose the wafer for 3s with Near-UV (405 nm) through our developed mask. The wafer was subsequently developed using MIF 726 (Hamatech-Steag wafer processor). After inspecting the wafer under a microscope, we subject it to a cycle of plasma descum to remove residual layers of photoresist (Anatech Resist Strip). We perform a Bosch fluorine process (Unaxis 770 Deep Silicon Etcher) to achieve a final etch depth of 5 $\mu$m, confirmed using the P-7 profilometer. We then strip the wafer of all resist using 2x10 minute wet bath soaks. Finally, we use the MVD-100 molecular vapor deposition system with the anti-stiction coating, 1H,1H,2H,2H-perfluorooctyltrichlorosilane (FOTS).

At this point, the wafer is ready to be used as a master mold of the pillar device. To make the microfluidic chips used for experiments, we mix polydimethylsiloxane (PDMS, Sylgard 184 Silicone Elastomer Kit, Dow Corning) in a ratio of 1:10, degas it in a desiccator, and pour it onto the master mold. We then bake it at 60 C° for 2 h. Devices were cut out from the wafer and we hole-punch 1.5mm-diameter holes at both inlets before binding them to a glass cover slip using an oxygen plasma oven.

**Microfluidic experiments and imaging**

We wet the device using LB medium with antibiotics immediately after the post-bake upon plasma binding the device to a clean glass bottom Petri-dish. We then inoculate 1 $\mu$L of the OD600 = 0.005 suspension in one inlet using a pipette tip. After inoculation, we seal both inlets using High-Vacuum Grease (Dow Corning). The sealed device is then submerged in Milli-Q water (preheated to 30 C°) to prevent the device from drying out and placed onto the microscope stage for imaging. We wait for ~2 h until the main arena of the device reaches sufficient cell density (10-50 cells in the field of view). During the time-lapse microscopy experiment, the temperature of the environment is maintained at 30 C°.

**Recording and quantifying bacterial motion**

Each device contain 3 levels of confinement for pillar regions and 4 levels of disorder for each level of confinement, amounting to a total of 12 unique pillar regions. Using a Nikon Eclipse Ti-E microscope equipped with a 20x Plan Fluor objective, mCherry fluorescence (49008, Chroma Inc.), and an Andor Neo sCMOS camera (Andor Technology plc.), we took video recordings for 40s (20 fps, 800 frames) at 12 regions with varying arrangements of pillars and one unconfined region without pillars. We repeat the process for a total of 7 times over the course of ~30 minutes to ensure sufficient cell trajectories.

We track between 68 and 605 cells for each region. To trace individual cells, we utilize the ImageJ plugin 'TrackMate' with sub-pixel precision (*35*). All tracks are inspected for quality control, and repetitive tracks, tracks that are not shown any sign of motion, and tracks appearing at the boundaries are removed. Raw tracking data $(x, y, t)$ is exported from ImageJ, and further processing and analysis is performed using Python scripts. All scripts and protocols used for analysis are available on Github (*34*). All raw images used for this research is publicly accessible in Dropbox: `https://www.dropbox.com/scl/fo/sfdxub50ny3niq647s0m1/AP2umhdJng3Pm1bxp02KQpc?rlkey=xad6wdjedgs7uwg8h2ty53sz0&st=9ozcb7om&dl=0`.

Starting from the raw tracking data, we assign each frame of a trajectory to a 'run' or 'tumble' state based on a combination of speed and turning angle thresholds. Speed is calculated as $v(t) = |\vec{\mathbf{v}}(t)| \equiv |\vec{\mathbf{r}}(t+\delta t) - \vec{\mathbf{r}}(t)| / \delta t$ and turning angle is defined as $\delta\theta(t) \equiv \tan^{-1}[\vec{\mathbf{v}}(t) \times \vec{\mathbf{v}}(t+\delta t) / \vec{\mathbf{v}}(t) \cdot \vec{\mathbf{v}}(t+\delta t)]$. We assign a frame as a run state when $v(t) \geq 12.4$ $\mu$m/s (half the unconfined mean speed) and $\delta\theta(t) \leq \pi/3$ (i.e., 60°). To identify tumble states, we choose the speed cutoff at half mean speed so cells tumble at times when $v(t) \leq 12.4$ $\mu$m/s.

##### Chord length calculation

We use an in-house Python script to simulate chord lengths in all pillar regions (Supplementary Fig. 8). Chord lengths are defined as randomly sampled, simulated straight paths from randomly positioned origins to the first intersecting pillar. To simulate chord lengths in each region, we recreate pillar arrangements for different levels of confinement and disorder. We then choose 400 randomly located origins from which 36 chords start; each chord ends at the first intersection with a pillar or edge of the device. A histogram of the resulting chords is used as an estimate of the probability density function of chord lengths in each region. Chord length distributions are not sensitive to the number of origins and the number of chords simulated.

##### Conditioned swim length distribution

Let $p$ be the probability density function (pdf) of swim lengths without confinement, and let $q$ be the pdf of chord lengths in a given environment. What is the expected swim length distribution if bacteria stopped when encountering a wall? If we independently draw one run length $r$ with probability density $p(r)$ and one chord length $s$ with probability density $q(s)$, the expected swim length in the environment is $z = \min(r, s)$. The pdf of the expected swim lengths is given by $f(z) = [1-Q(z)]\, p(z) + [1-P(z)]\, q(z)$, where $Q$ and $P$ are the cumulative distribution functions of chord lengths and free-space swim lengths, respectively. The conditioned swim length distributions shown in Fig. 6 are calculated from the empirically observed swim length distribution and the numerically determined chord length distributions using the formula above.

**Mean-square displacement (MSD) calculation**

To compute mean-square displacements (MSDs), we divide tracks into non-overlapping time segments ($dt$ = 0.05 s, corresponding to one frame) as $MSD(\tau) = \left\langle (r\,(t+\tau) - r\,(t))^2 \right\rangle$, where $\tau$ is the lag time. The range of $\tau$ is chosen to ensure that the minimum number of trajectories is at least 50, with a maximum $\tau$ of 20 seconds. The ensemble-averaged MSDs (eMSDs) take the average over bacterial trajectories for a given region. All time- and ensemble-averaged MSDs are shown in Supplementary Figs. 4 and 5, respectively. To fit eMSDs at early lag times ($\tau$ ¡ 0.5 s), we use a power law, $y = \beta x^{\alpha}$ where $\beta$ and the exponent $\alpha$ are two parameters to adjust so as to produce the best fit, using the Python package Scipy (*36*). We present a heatmap to show how the exponent $\alpha$ varies across conditions.

**Swim and stall simulations**

We simulate stochastic run and tumble dynamics in two dimensions via an in-house Python script. Tumbles are initiated at a rate $T_{\text{freq}}$ per second; each tumble duration is drawn from an exponential distribution with mean $\langle T_{\text{tumble}} \rangle = 0.2$s (Fig. 2D). Tumble initiations are independent at each time point for each simulated bacterium. This results in exponentially-distributed run duration with mean $\langle T_{\text{run}} \rangle = 1/T_{\text{freq}}$. At time points corresponding to the end of a tumble, another tumble is immediately initiated (corresponding to a failed reorientation) with probability $T_{\text{fail}}$. Simulations are performed at our experimental resolution, with $dt = 0.05$s for $n = 200$ bacteria over a 60 second interval. MSDs for simulated trajectories are calculated as for experimental trajectories.

#### Supplementary Figures

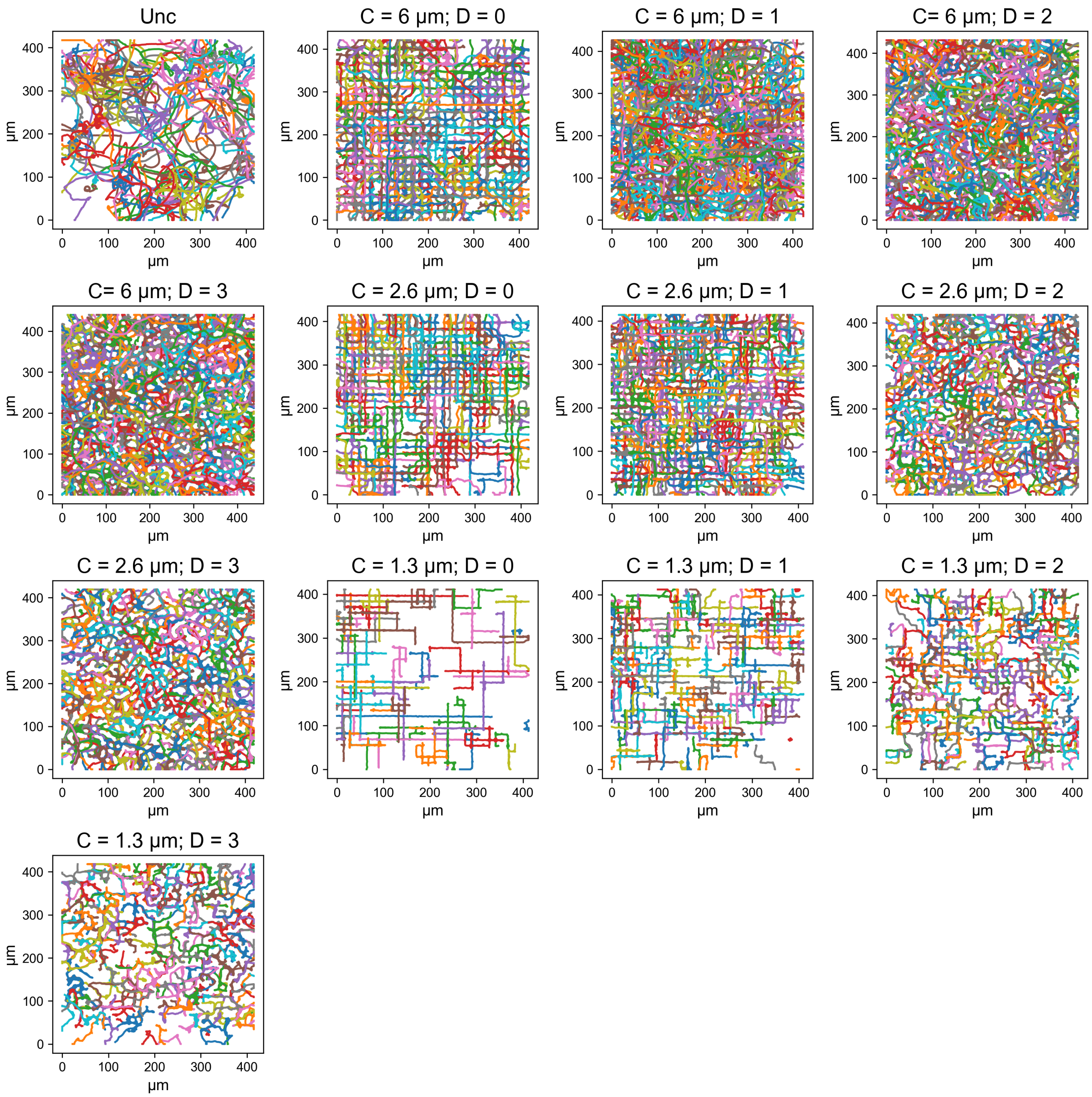


**Figure S1**: **Bacterial track overlays of all regions examined.** The label 'unc' denotes the condition 'unconfined'. The colors of the trajectories are used to make bacterial tracks distinguishable.

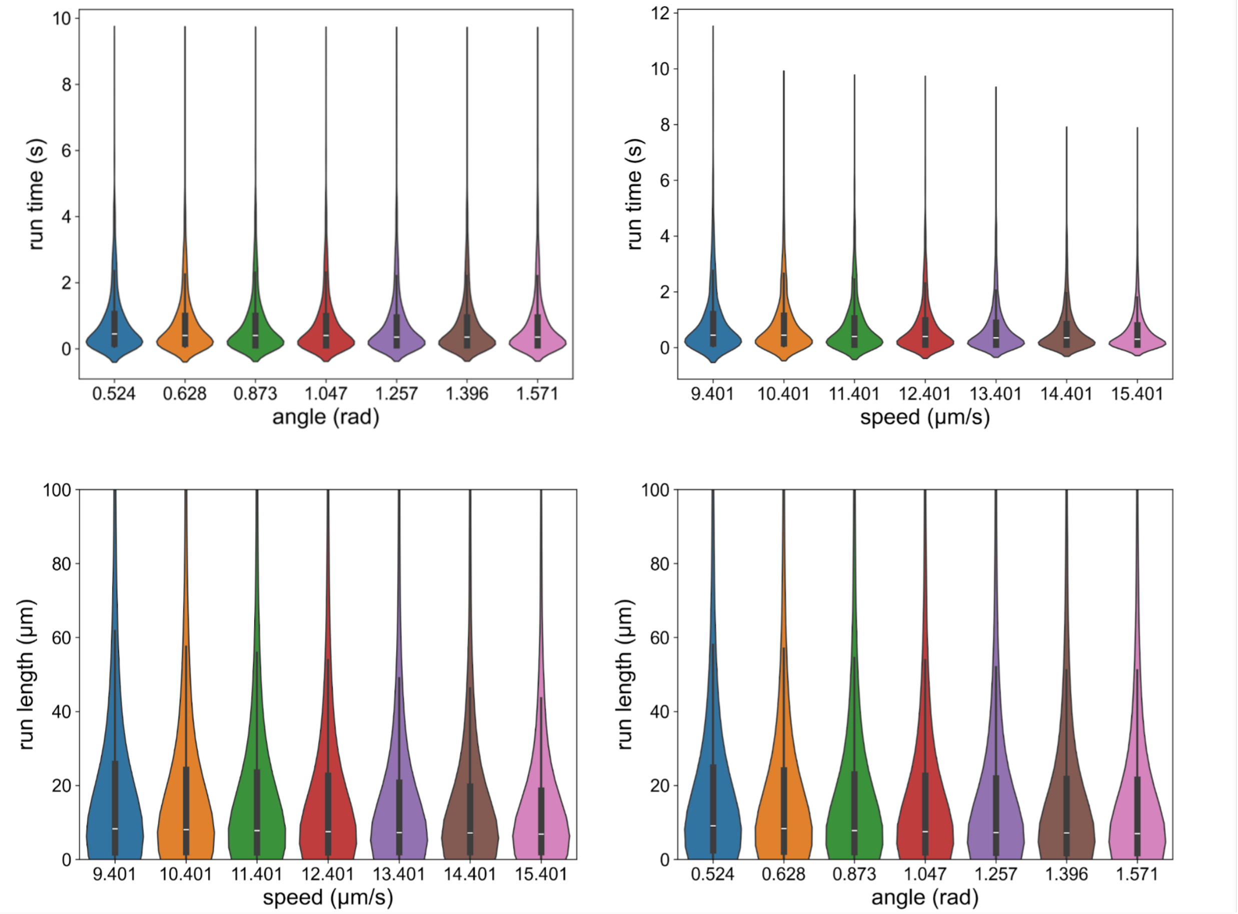


**Figure S2**: **Sensitivity checks of speed and angle thresholds.** The violin plots show the distributions of run times and run lengths across different speed and angle thresholds. The distributions are not sensitive to various thresholds tested.

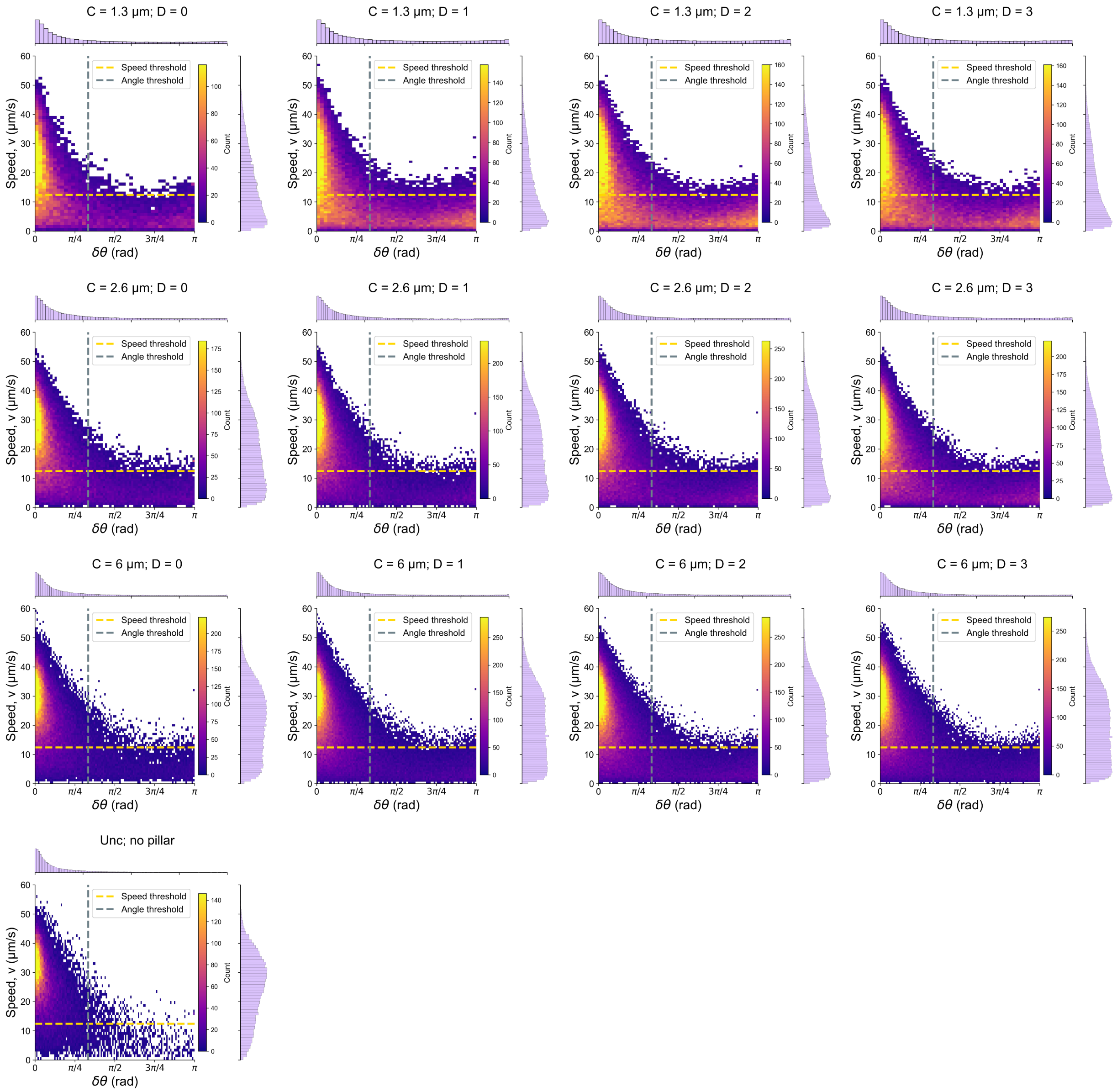


**Figure S3**: **Speed-angle jointplot of all regions.** The yellow dash line denotes speed threshold, and the gray dash line denotes velocity orientation angle threshold.

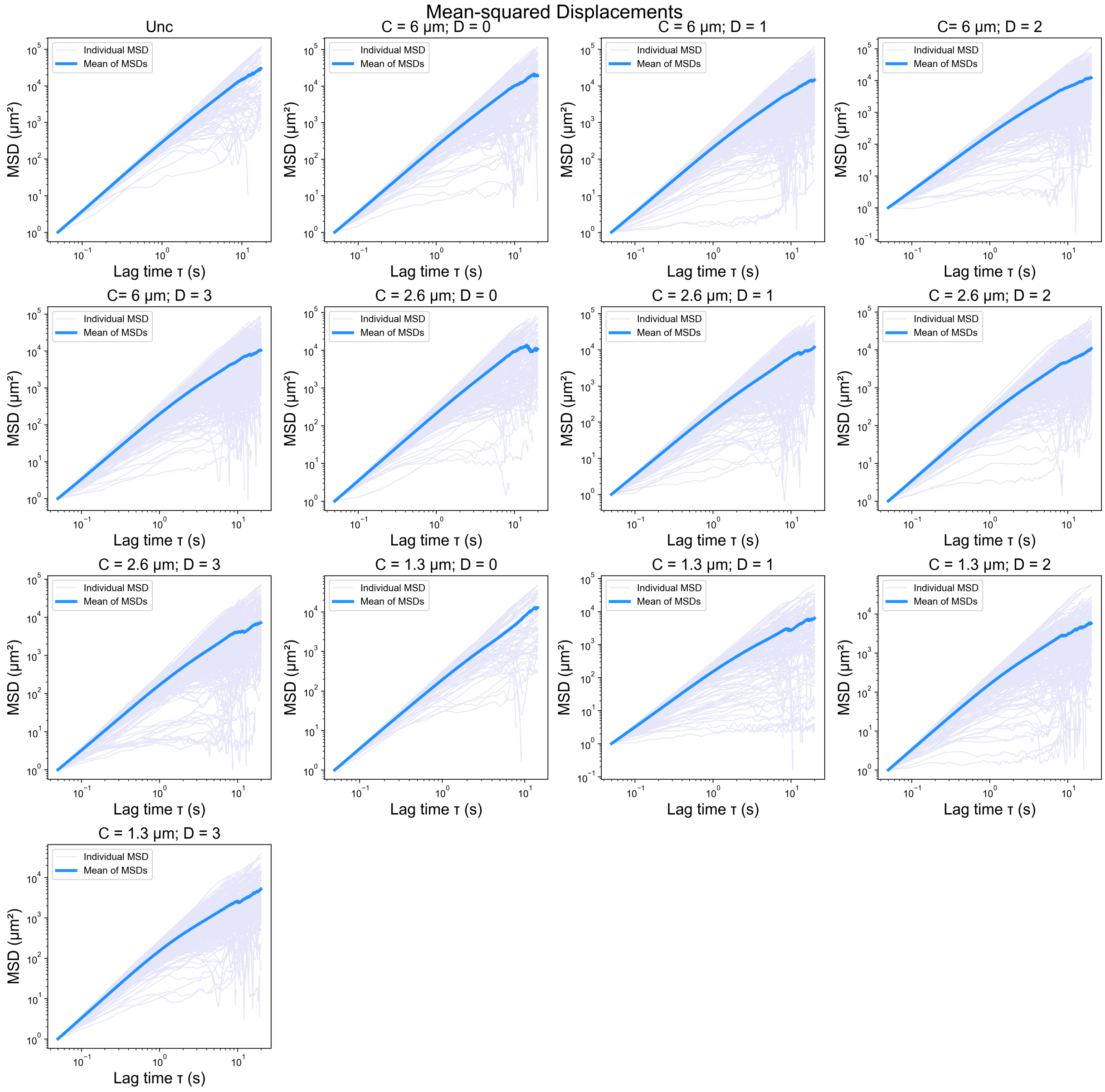


**Figure S4**: **MSD plots under all experimental conditions.** Individual MSDs of all regions are labeled in purple. The ensemble-average MSD of a region is labeled in blue.

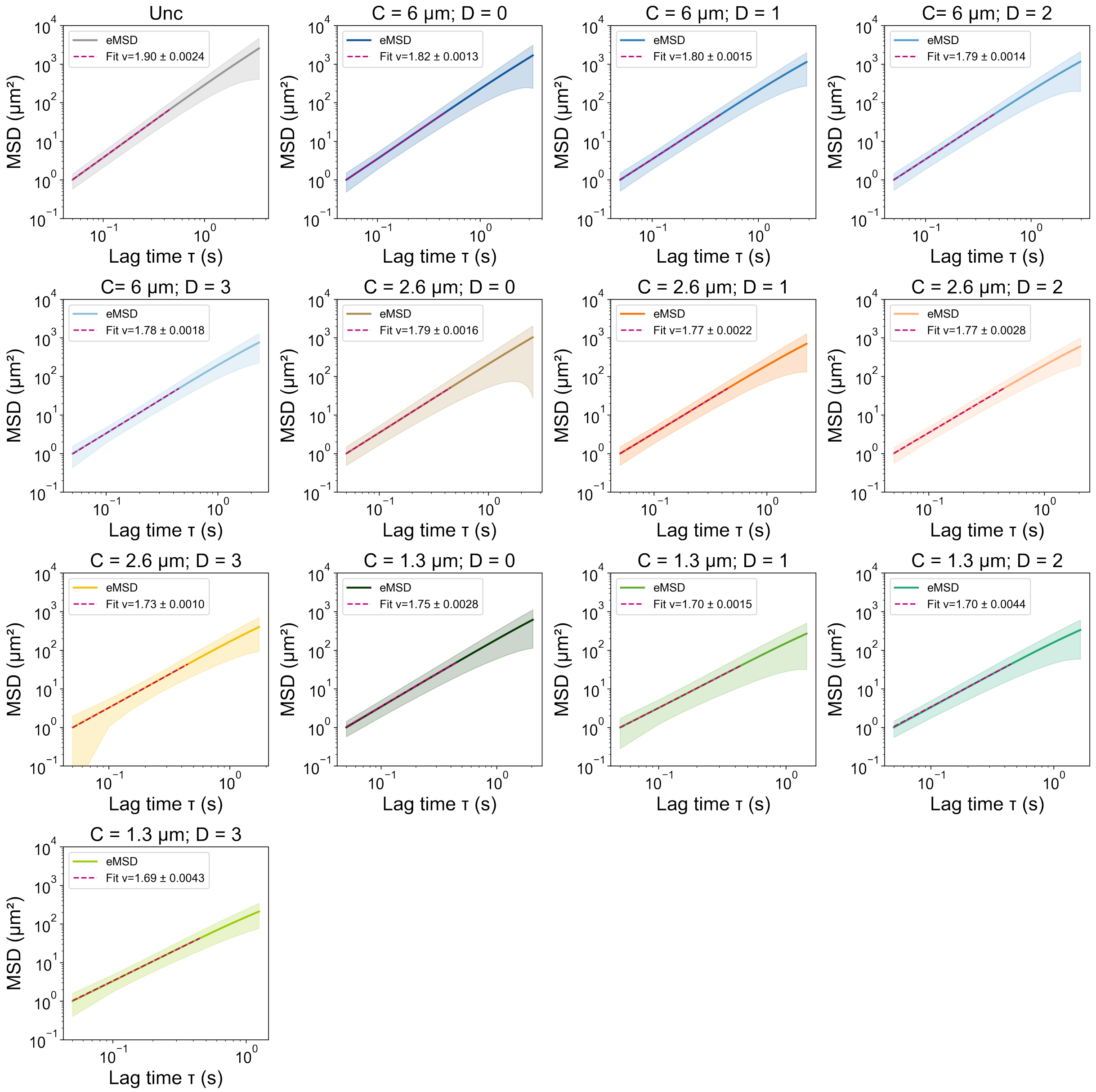


**Figure S5**: **Ensemble-average MSDs (eMSDs) and power-law fitting curves at early lag times for all conditions.** The bipolar bands indicate standard deviations (eMSDs ± std). The fit for early lag times are labeled in pink dashed lines by *Scipy* package.

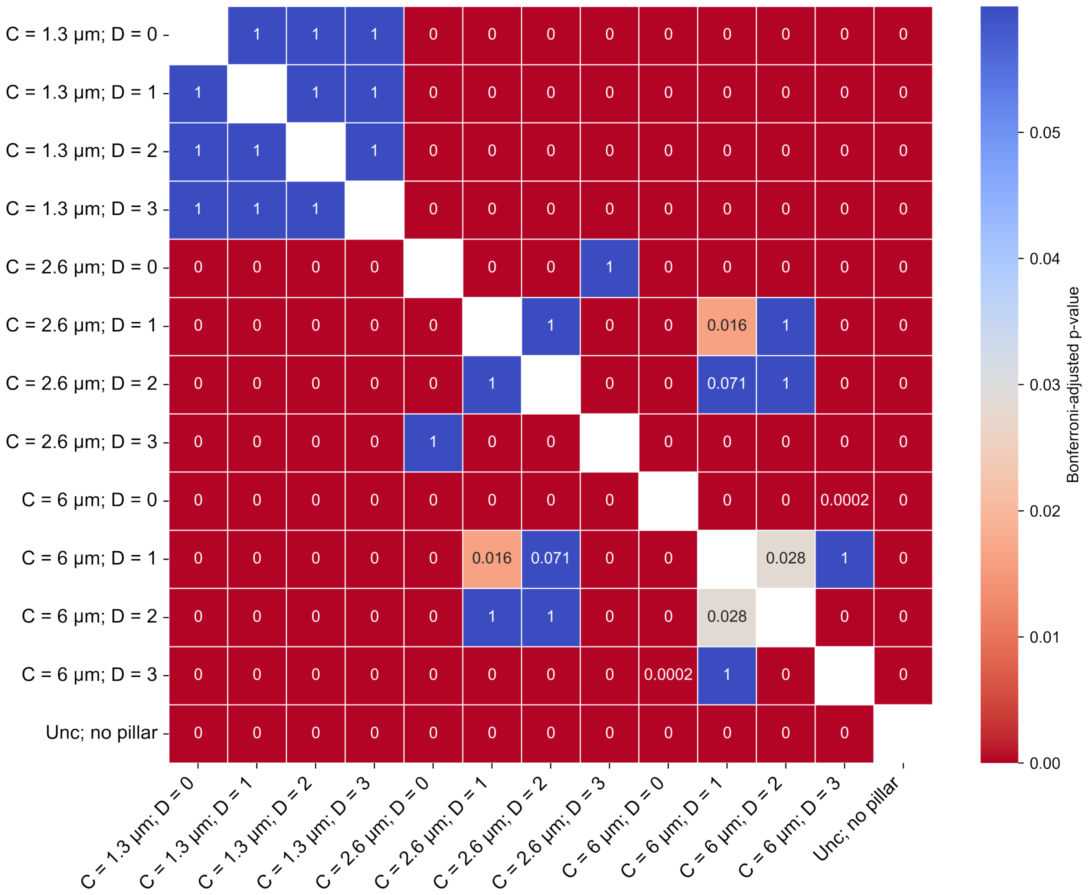


**Figure S6**: **Pairwise Mann-Whitney U test of swim durations between conditions.** P values are corrected by the Bonferroni method. The significance value is indicated by a color spectrum (red: very significant; gray: significant; blue: not significant).

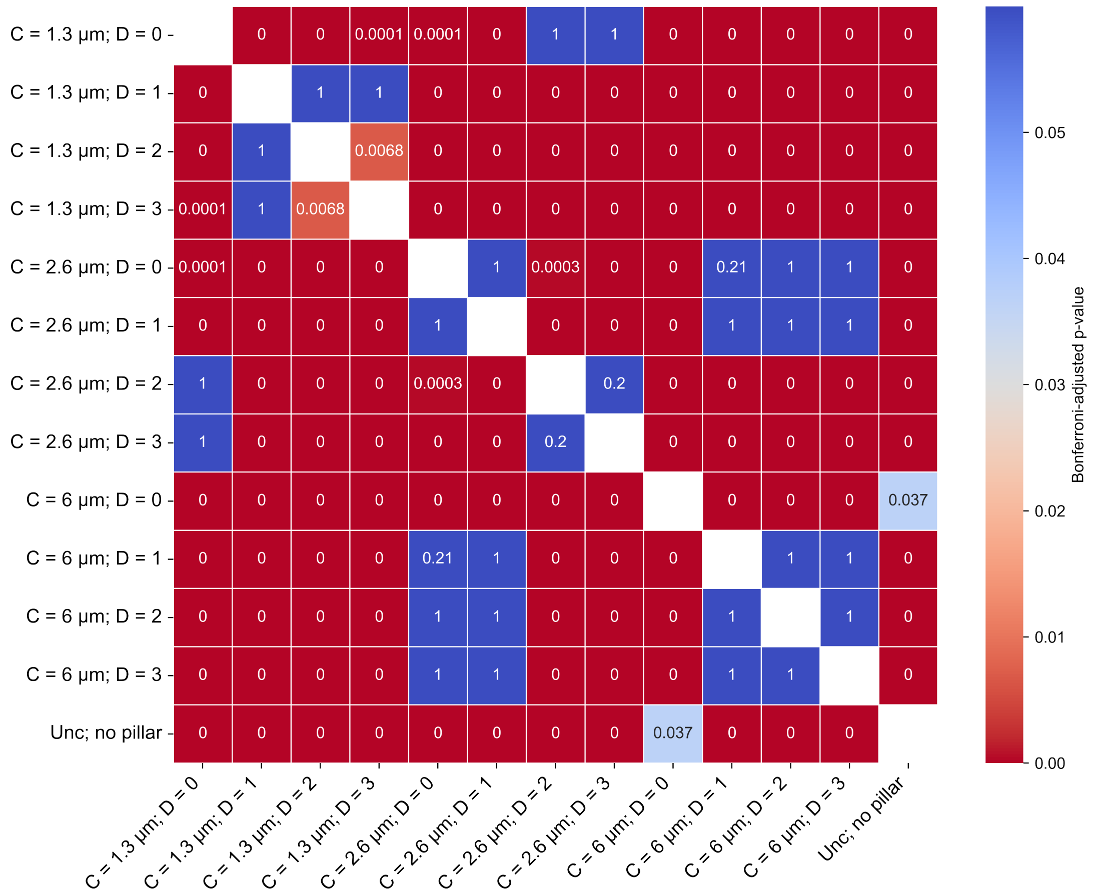


**Figure S7**: **Pairwise Mann-Whitney U test of stall durations between conditions.** P values are corrected by the Bonferroni method. The significance value is indicated by a color spectrum (red: very significant; gray: significant; blue: not significant).

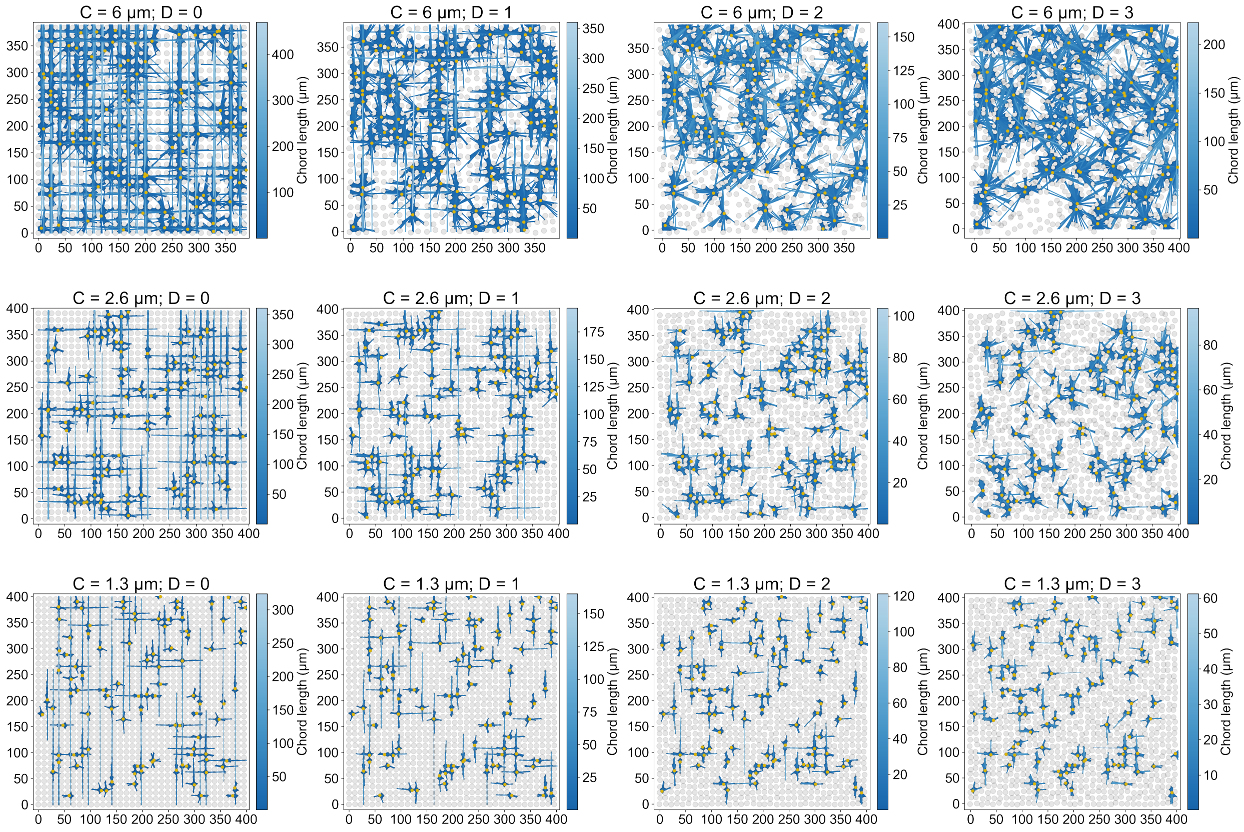


**Figure S8**: **Chord length simulations of all regions.** Every simulated region has one hundred origins indicated by yellow circles and each origin has one hundred chords. The color bars show the range of chord lengths.

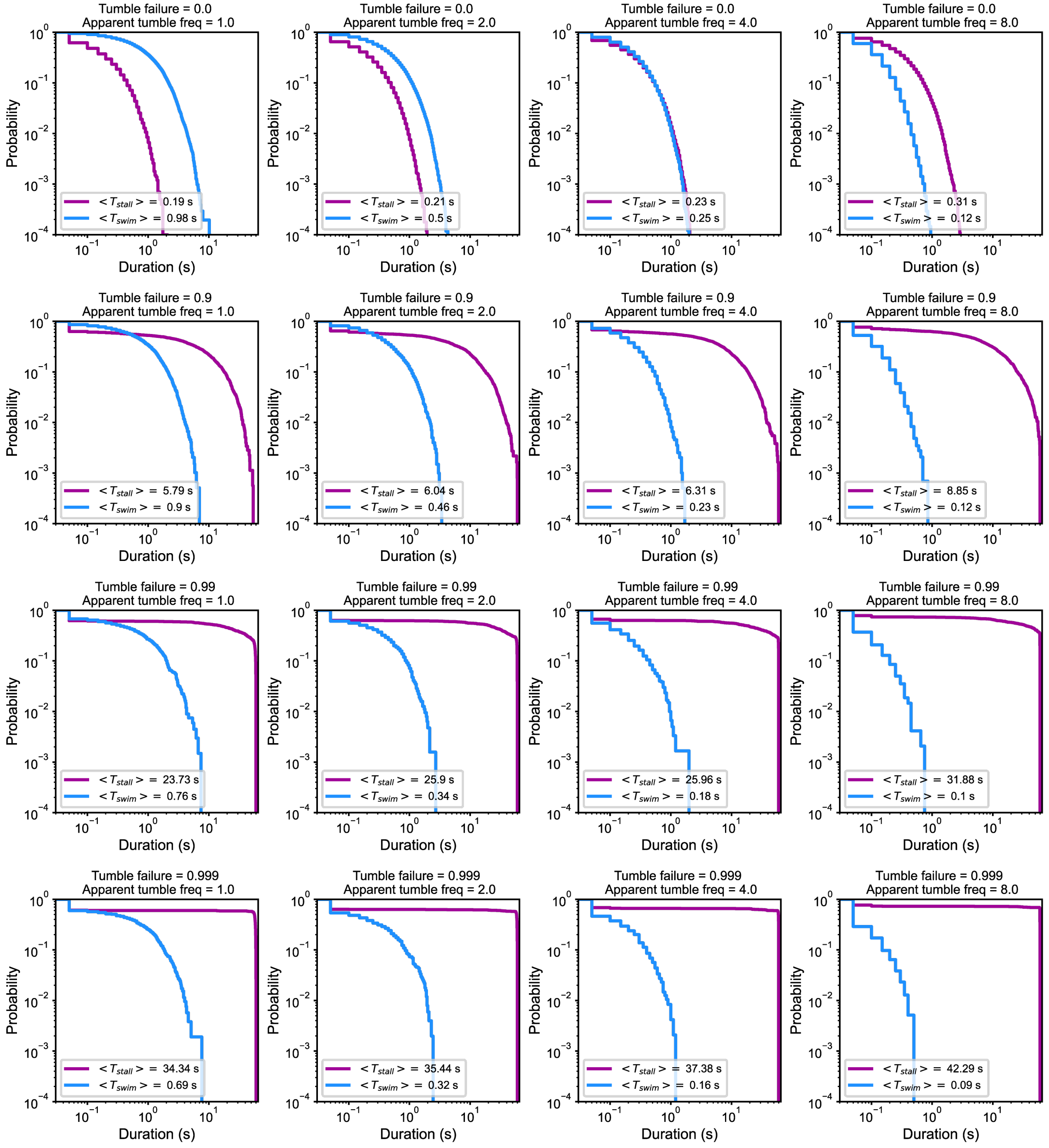


**Figure S9**: **Complementary cumulative distribution function (CCDF) plots of stall and swim durations under all simulated conditions.** In each subplot, CCDF for stall duration is labeled in magenta and CCDF for swim duration is labeled in blue. Each subplot has mean stall and swim durations indicated in the inset.

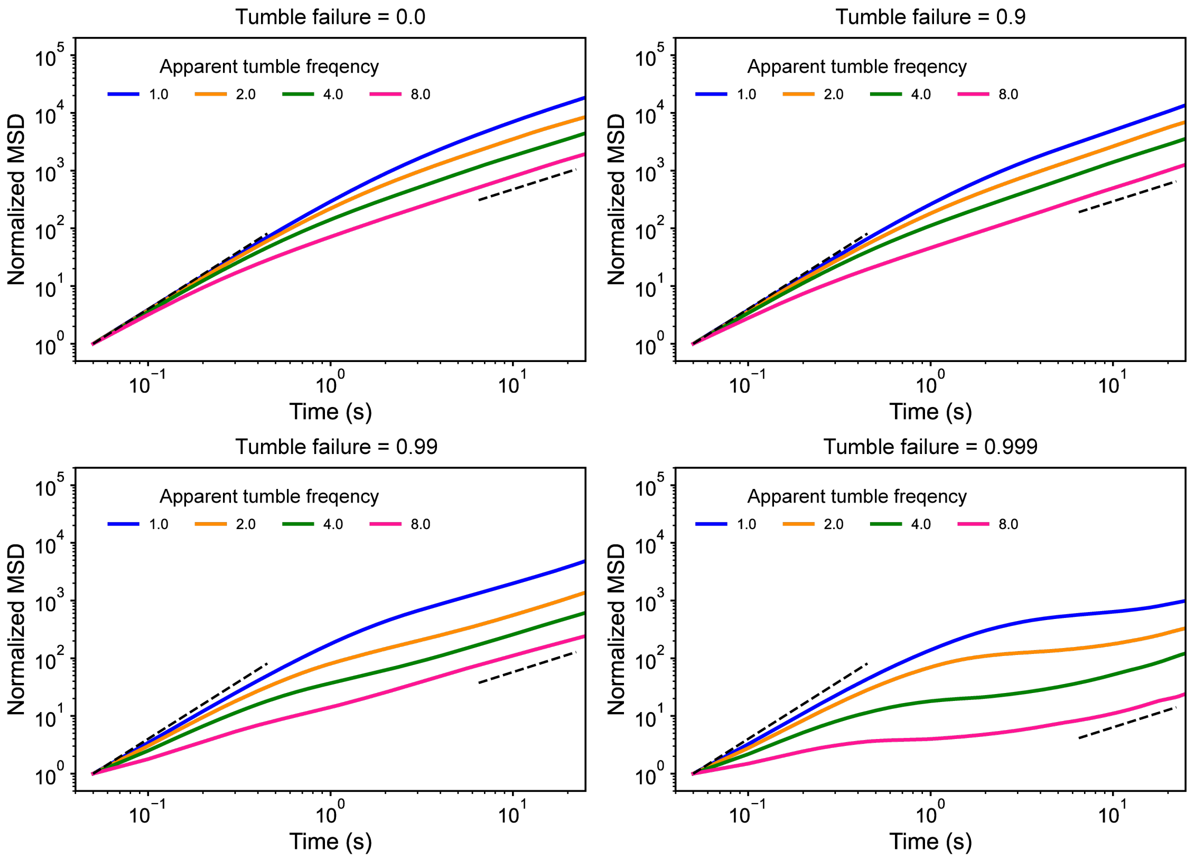


**Figure S10**: **Ensemble-average MSDs varying apparent tumble frequency.** Each subplot has a fixed tumble failure probability: 0, 0.9, 0.99, and 0.999, respectively.

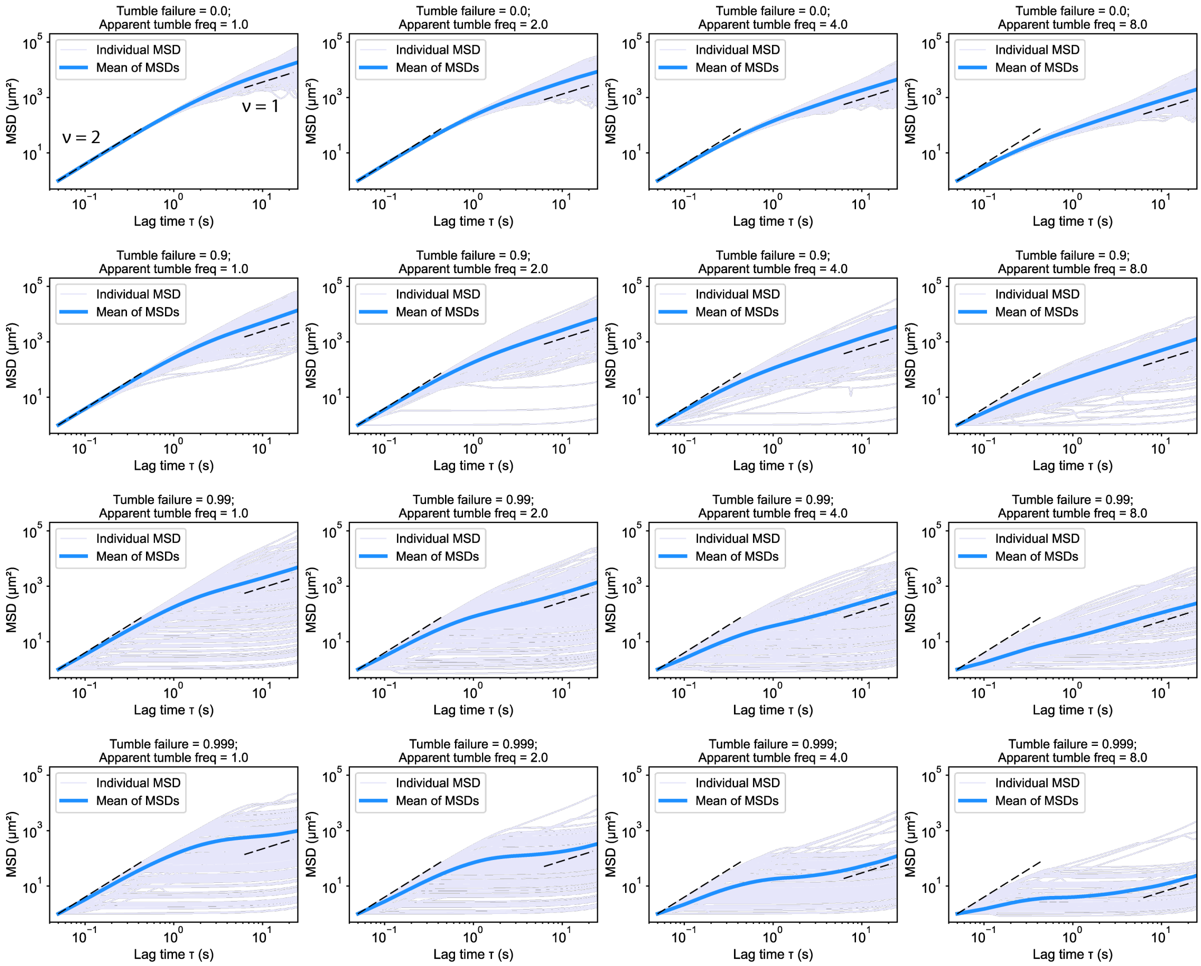


**Figure S11**: **Simulated MSD plots of all conditions.** Individual MSDs of all regions are labeled in purple. The ensemble-average MSD is labeled in blue in a subplot.

#### Supplementary Movies

**Caption for Movie S1. Fluorescently labeled *E. coli* HCB 33 explore the microfluidic region C = 6 $\mu m$, D = 0 in real time.** The frame rate is 20 frames per second.

**Caption for Movie S2. Fluorescently labeled *E. coli* HCB 33 explore the microfluidic region C = 2.6 $\mu m$, D = 0 in real time.** The frame rate is 20 frames per second.

**Caption for Movie S3. Fluorescently labeled *E. coli* HCB 33 explore the microfluidic region C = 1.3 $\mu m$, D = 0 in real time.** The frame rate is 20 frames per second.
